# Meta-analytic microbiome target discovery for immune checkpoint inhibitor response in advanced melanoma

**DOI:** 10.1101/2025.03.21.644637

**Authors:** Xinyang Zhang, Himel Mallick, Ali Rahnavard

## Abstract

Immune checkpoint inhibitors (ICIs) have revolutionized melanoma treatment, yet patient responses remain highly variable, underscoring the need for predictive biomarkers. Emerging evidence suggests that gut microbiome composition influences ICI efficacy, though findings remain inconsistent across studies. Here, we present a meta-analysis of seven melanoma-associated microbiome cohorts (N=678) using a standardized computational pipeline to integrate microbial species, biosynthetic gene clusters (BGCs), and functional pathways. We identify *Faecalibacterium* SGB15346 as a key species enriched in responders, alongside RiPP biosynthetic class and pathways involved in short-chain fatty acid fermentation. Conversely, dTDP-sugar biosynthesis correlates with non-response. Our results highlight microbial signatures and metabolic pathways associated with ICI outcomes, offering potential targets for microbiome-based interventions in personalized immunotherapy.

## Main

Advancements in cancer immunotherapy have significantly improved survival rates for melanoma patients, particularly through the use of immune checkpoint inhibitors (ICIs) targeting programmed cell death protein 1 (PD-1) and cytotoxic T-lymphocyte-associated protein 4 (CTLA-4)^1–4^. However, clinical responses to ICIs remain inconsistent, suggesting that factors beyond tumor-intrinsic properties and host immune responses influence therapeutic efficacy^2,5^. Increasingly, research highlights the role of the gut microbiome in modulating immune responses, with specific microbial taxa and functional pathways associated with treatment outcomes^6,7^. Understanding these interactions is critical for developing microbiome-based strategies to enhance immunotherapy efficacy.

Early studies indicate that specific gut microbial taxa, such as *Akkermansia muciniphila, Faecalibacterium prausnitzii, Bifidobacterium longum*, and *Bacteroides caccae*, are enriched in ICI responders^5,8–10^. Additionally, fecal microbiota transplantation (FMT) from responders into non-responders can restore PD-1 blockade sensitivity, underscoring the causative role of the microbiome in immunomodulation^11,12^. Dietary factors have also emerged as a promising avenue, with higher fiber intake correlating with improved progression-free survival in melanoma patients treated with immunotherapy^13^.

Despite promising findings, research on microbiome-ICI interactions has produced inconsistent results, primarily due to variability in study design, sample collection methods, dietary influences, medication use, and microbiome sequencing techniques^6,14^. Meta-analysis serves as a powerful tool to overcome these challenges, integrating data across multiple studies to enhance statistical power and uncover robust microbial signatures associated with ICI response. However, previous meta-analyses have been limited in scope, largely focusing on microbial taxa while overlooking biosynthetic gene clusters (BGCs) and metabolic pathways that may play critical roles in immunotherapy outcomes^14–16^. Moreover, these studies have analyzed a smaller number of cohorts, restricting their ability to achieve cross-study reproducibility and draw generalizable conclusions. Additionally, prior meta-analyses have not utilized the most up-to-date profiling tools, potentially missing key microbial signatures and functional elements relevant to ICI response.

Recognizing these limitations, we compiled publicly available whole metagenome shotgun sequencing (MGS) datasets of gut microbiomes from melanoma patients receiving immunotherapy and conducted a comprehensive meta-analysis across seven studies (N = 678). This analysis integrates multiple layers of biological information—including taxonomic profiles, biosynthetic gene clusters (BGCs), and functional pathways—to examine differences in the microbiota’s compositional and functional characteristics between responders (R) and non-responders (NR), providing a more holistic view of microbiome-ICI interactions (**Fig. 1**). Unlike previous taxonomy-focused studies, which relied on MetaPhlAn 2^17^, we utilize MetaPhlAn 4^18^, which represents a substantial expansion with a significantly larger reference database, improved taxonomic resolution, and enhanced detection of previously uncharacterized species. This allowed us to more accurately and comprehensively connect actionable microbial features with immunotherapy response, laying the foundation for microbiome-based multimodal therapeutic strategies in melanoma treatment.

**Figure 1:**
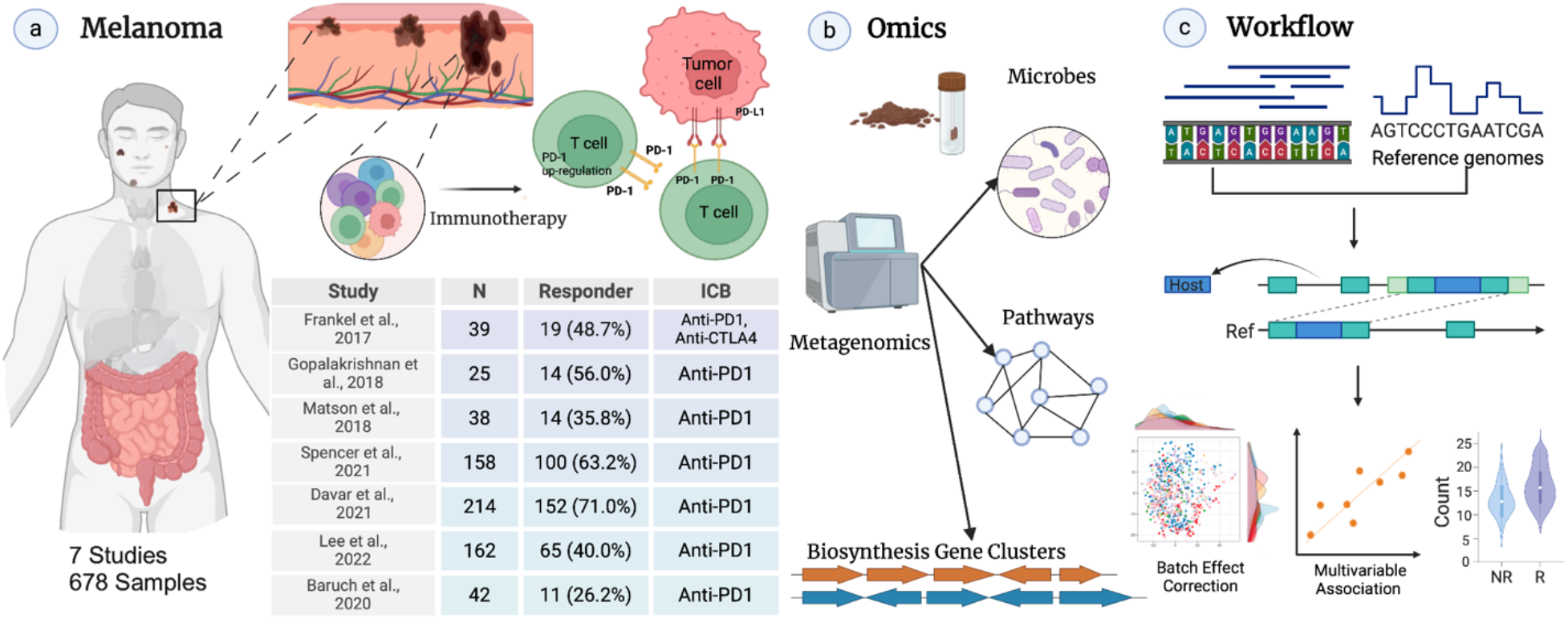
Overview of Study Design, Omics Analysis, and Workflow. **a**, Melanoma progression and immune response: Melanoma advances through skin layers, involving the immune response between T cells and tumor cells. Immunotherapy targeting PD-1/PD-L1 enhances T cell activity against tumors. Data include 678 samples from 7 studies. **b**, Multi-Omic Data Integration: Microbial communities, metagenomics, and pathway analyses reveal biosynthesis gene clusters and microbial pathways influencing melanoma progression and immunotherapy response. **c**, Computational Workflow: Microbial genomic data undergoes host genome removal before alignment with reference genomes. Batch effect correction is applied, and multivariable models assess associations between microbial features and immunotherapy outcomes, distinguishing responders (R) from non-responders (NR).

## Results

### Clarifying Variance: Beta Diversity and Clustering & Batch Effect Correction

We identified seven studies that met the inclusion criteria, which required a focus on melanoma patients undergoing ICI therapy targeting PD-1 and/or CTLA-4, the use of gut microbiome profiling through shotgun metagenomics sequencing, and the availability of clinical response data stratified by responders and non-responders. Additionally, studies had to provide publicly accessible or author-shared microbiome data with taxonomic and/or functional annotations while employing standardized or comparable DNA extraction and sequencing techniques to minimize methodological bias. A minimum cohort size of at least 10 patients per response group was also required to ensure statistical validity. Despite meeting these criteria, the included studies exhibited variability in DNA extraction toolkits, which can significantly impact downstream analyses due to differences in DNA yield^19,20^.

To assess inter-study variability, we examined beta diversity, which measures differences in microbial composition between samples and evaluated the similarity of microbial profiles across studies (**Methods**). Variability in microbial composition can arise from differences in patient populations, sample processing methods, sequencing techniques, or underlying biological heterogeneity, all of which can introduce batch effects that obscure true biological signals. By analyzing beta diversity metrics, we aimed to determine the extent of divergence between studies and identify clusters of studies with similar microbial communities. We found that studies such as ‘Davar,’ ‘Spencer,’ ‘Gop,’ and ‘Matson’ cluster closely together, suggesting high similarity in their microbial composition. At the same time, ‘Frankel,’ ‘Lee,’ and ‘Baruch’ form a separate group, indicative of greater dissimilarity (**Fig. 2a(Left)**).

**Figure 2:**
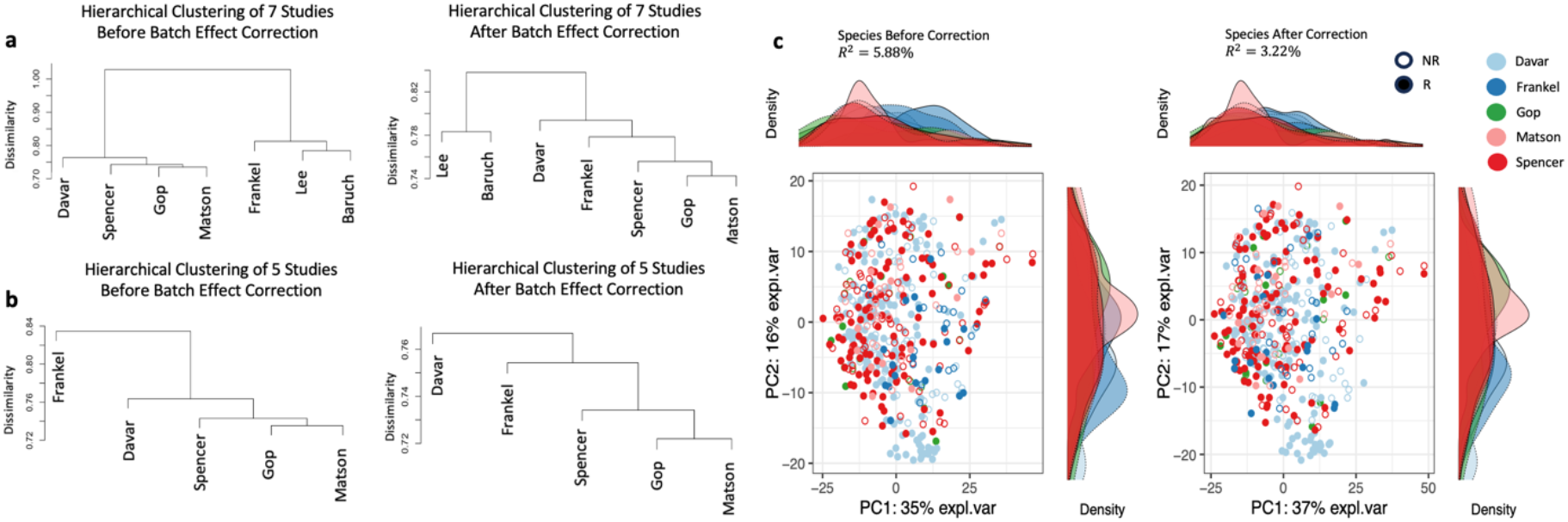
Beta Diversity Clustering and the Effect of Batch Correction. **a**, Hierarchical clustering of seven studies before(left) and after(right) batch effect correction, respectively. Before batch effect correction, distinct groupings influenced by batch effects were shown. The clustering pattern suggests that study-specific biases contribute significantly to the observed dissimilarities. After batch effect correction, clustering of the same studies after batch effect correction reveals a reduction in batch effects, with studies clustering more closely together. However, some degree of dissimilarity remains, suggesting that while the correction mitigates batch effects, it does not entirely eliminate them. Residual batch effects may still influence clustering patterns, likely due to inherent study-specific differences that persist despite correction. **b**, Conducting the same analysis for a subset of five studies, before batch correction, datasets cluster primarily by study, indicating strong batch effects. After correction, batch effects are reduced, but complete removal was not achieved. Notably, despite batch correction(shown in a), the seven studies did not fully integrate into a single branch, leading us to refine the dataset selection. Considering factors such as differences in DNA extraction kits and sequencing protocols, we focused on a subset of five studies, which, after batch correction, successfully clustered under the same major branch. **c**, display Principal Component Analysis (PCA) with Density plots per component^21^ of species-level diversity. Pre-correction (Left), the data points cluster by study batch, showing greater batch-related variation. Post-correction (Right), batch effects are reduced, and the studies align more closely while still retaining meaningful biological signals.

The hierarchical clustering of the seven studies after batch effect correction demonstrates a reduction in inter-study dissimilarity, as evidenced by lower dissimilarity values (**Fig. 2a(Right)**). However, some level of study-specific clustering remains, with studies such as ‘Lee’ and ‘Baruch’ forming distinct clusters separate from others. This residual clustering suggests that while batch effect correction successfully mitigated a substantial portion of the inter-study variability, it did not eliminate all batch effects. The persistence of these effects may reflect unmeasured confounders, experimental differences, or inherent biological variability between studies. These findings highlight the challenges of completely normalizing batch effects in complex datasets, underscoring the need to interpret inter-study comparisons even after correction.

The differences in microbial community profiles across studies emphasized the need for subsequent batch (study) effect correction to mitigate potential inter-study variability. To this end, we applied the MMUPHin workflow and quantified the variance explained by batch effects by permutational multivariate analysis of variance (PERMANOVA). For the refined dataset of five studies for species, PERMANOVA results before batch effect correction (**Fig. 2b**) showed that dataset grouping explained 5.88% of the variance (R^2^ = 0.0588, p = 0.001), indicating a moderate influence of study-specific effects. After batch effect correction, the variance explained by dataset grouping was reduced to 3.22% (R^2^ = 0.0322, p = 0.001). This reduction demonstrates that batch correction effectively minimized inter-study variability, though a small residual effect remains. The increase in residual variance (from 94.1% to 96.8%) further confirms that most of the variability is now attributed to biological or experimental noise rather than systematic batch effects. This reduction in cohort-driven variability highlights the success of the batch correction procedure in minimizing study-specific biases, allowing for a more accurate assessment of the underlying biological signals.

### Faecalibacterium SGB15346 is strongly linked to responders to ICI

For species-level taxonomic profiling, we included five out of seven studies that exhibited high taxonomic similarity, ensuring robust comparisons among datasets. A total of 474 metagenomes from five independent cohorts, comprising 299 non-responder samples and 175 responder samples collected between 2017 and 2021, were analyzed. To identify gut microbiome features associated with immunotherapy outcomes, taxonomic profiling was performed for each cohort, followed by integration across studies with batch effect correction. The Compound Poisson Linear Model (CPLM) was then applied to link microbiome features to response status (responder vs. non-responder). Species-level microbiome features were selected based on their significance across all five studies (p < 0.05) and low-to-moderate heterogeneity (*I*^2^< 60%).

Significant species were ranked by their coefficients (**Fig. 3a**), with positive and negative associations distributed symmetrically. Among species positively correlated with responders, *GGB9534 SGB14937* and *Faecalibacterium SGB15346* showed the strongest associations. Notably, *Faecalibacterium SGB15346* emerged as the most significant species associated with responders (p < 0.01, coefficient = 0.80, *I*^2^= 0, and q-val.fdr< 0.05).

**Figure 3:**
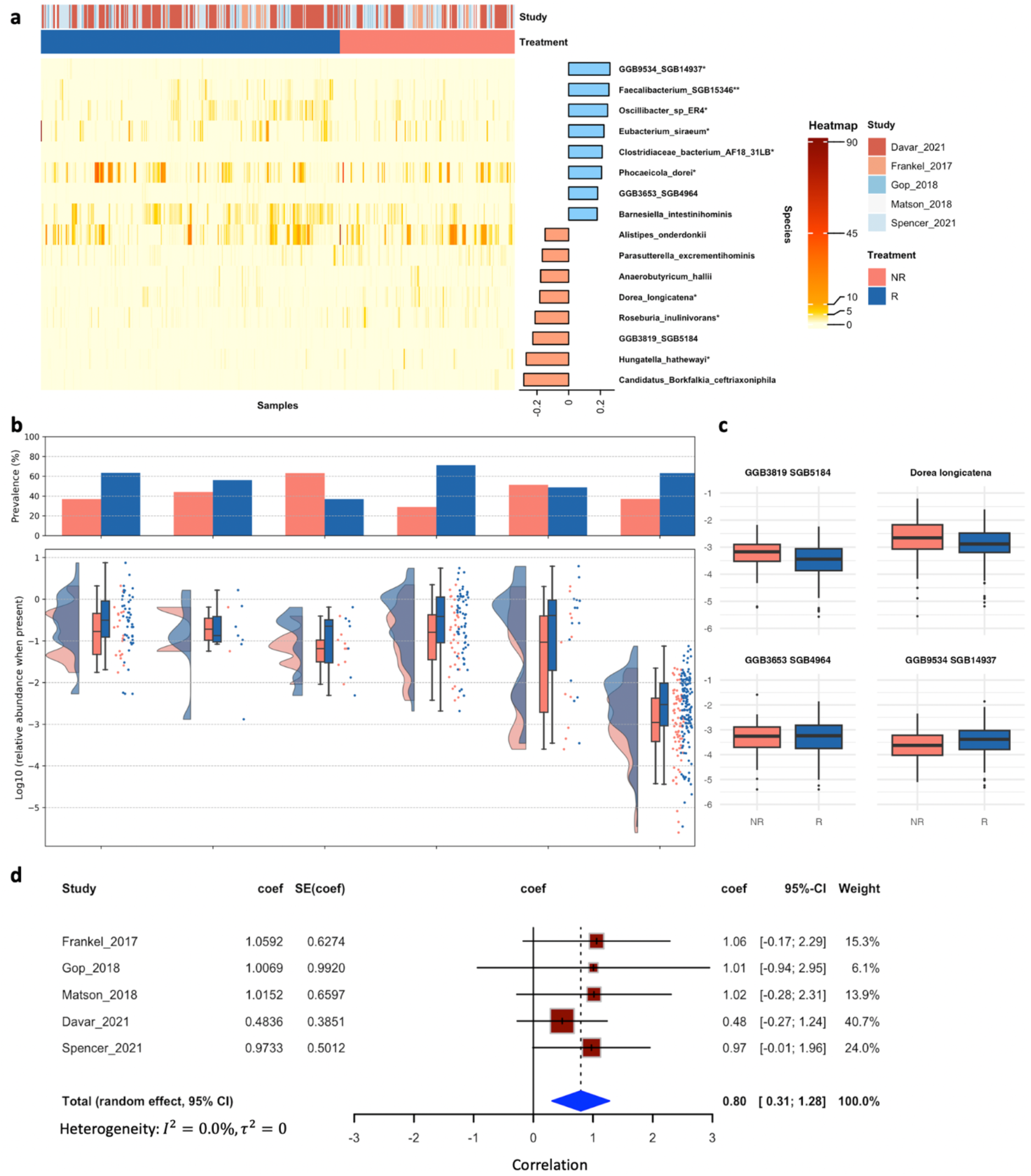
Microbial species associated with immunotherapy response in melanoma. **a**, Heatmap of significant microbial species across studies, grouped by response status (response (R) and non-response (NR)) to immunotherapy across multiple melanoma studies. Rows represent microbial species, and columns correspond to individual samples, organized by study and treatment response. The gradient color scale reflects species abundance, with red hues indicating higher levels. Notable species, such as *Faecalibacterium SGB15346* and *GGB9534 SGB14937*, positively correlate with treatment response, whereas others, like *Dorea longicatena*, exhibit differential associations. The accompanying bar graph on the right displays the strength of these associations, represented as log2 fold changes. Species names marked with an asterisk (*) denote statistically significant associations. **b**, The raincloud plot depicts the prevalence and relative abundance of *Faecalibacterium* SGB15346 across five studies, highlighting its consistent enrichment in responders. The top panel shows the prevalence (percentage of samples where the species is detected) in each study, with red and blue bars representing different conditions. The bottom panel presents the log10-transformed relative abundance of microbial species when present, visualized using boxplots and violin plots. The distribution of data points highlights differences in microbial composition between responders and non-responders. **c**, Boxplots compare the relative abundances of four key species (*Dorea longicatena, GGB3653 SGB4964, GGB3819 SGB5184*, and *GGB9534 SGB14937*) between responders (R) and non-responders (NR). Differences in microbial composition confirm significant variations between groups, suggesting that specific gut microbiota may play a critical role in influencing immunotherapy outcomes in melanoma. **d**, Forest plot of the meta-analysis for the association between the abundance of *Faecalibacterium SGB15346* and response to treatment. Each horizontal line represents an individual study, with the corresponding effect size (coef) and 95% confidence interval (CI) shown. The red squares represent the point estimates of effect sizes, with their sizes proportional to the weight of the study in the meta-analysis. The blue diamond at the bottom represents the pooled effect size from the random-effects model (coef = 0.80, 95% CI: [0.31; 1.28]). The dashed vertical line indicates no effect (coef = 0). Heterogeneity across studies is low (I^2^ = 0.0%, τ ^2^ = 0).

To further investigate the association between gut microbiome features and immunotherapy response, we examined the prevalence and relative abundance of *Faecalibacterium SGB15346*, the most strongly correlated species with responders (**Fig. 3b**). The bar chart in the upper panel shows the prevalence of this species across individual cohorts, with a general trend of higher prevalence in responders. However, the patterns within individual studies are not always consistent or clear due to variability in sample size and cohort-specific factors. Similarly, the violin plots in the lower panel, which depict the log10-transformed relative abundance, show considerable overlap between responders and non-responders within some individual studies.

Notably, when data from all five studies are combined, a clear and statistically significant pattern emerges, with responders showing a higher prevalence and relative abundance of *Faecalibacterium SGB15346* (FDR < 0.05 in the combined analysis). This highlights the power and beauty of meta-analysis: by aggregating data across multiple studies, we can overcome individual study limitations and achieve greater statistical robustness, uncovering patterns that might otherwise remain obscured. The consistency of this association across the combined dataset underscores the potential of *Faecalibacterium SGB15346* as a robust biomarker for immunotherapy response.

Taken together, these results reveal key gut microbiome features that distinguish responders from non-responders. *Faecalibacterium SGB15346*, in particular, emerged as the most significant species, suggesting its potential as a diagnostic biomarker or therapeutic target. The consistent trends observed across multiple cohorts and analytical approaches underscore the robustness of these findings. Future work should focus on uncovering the mechanisms by which these microbiome features influence immune modulation and therapy outcomes in melanoma patients.

### Composition of the pathway analysis and biosynthesis gene cluster analysis

To provide additional functional context, we integrated pathway abundance data with gene ontology (GO) terms^22^. This analysis linked specific pathways to biological processes such as immune cell activation, response to cytokines, and cellular stress responses. The integration of pathway data with GO terms enriched our understanding of how these pathways contribute to therapeutic success. The combined analysis of gene families and pathways reveals a coordinated network of immune-related mechanisms driving the effectiveness of cancer immunotherapy. This included activating specific gene families and pathways that work together to enhance immune surveillance and tumor suppression. These revealed an array of gut-associated microbial pathways that positively correlated with response compared with the non-response samples (**Fig. 4a**). These included GABA, 4-aminobutanoate degradation V (pathway 5022). GABA inhibits electrical activity in melanoma/keratinocyte cocultures^23–25^. Cancer cells exhibit increased glucose uptake to meet their high metabolic demands, driving glycolysis and glucose-derived biosynthetic pathways, including dTDP-α-D-ravidosamine and dTDP-4-acetyl-α-D-ravidosamine biosynthesis (pathway 7688) ^26,27^. The overactivity of these pathways, which are involved in glycosylation, may negatively impact anti-PD-1 therapy efficacy by promoting tumor immune evasion or metabolic adaptation, as suggested by their negative correlation with responders.

**Figure 4:**
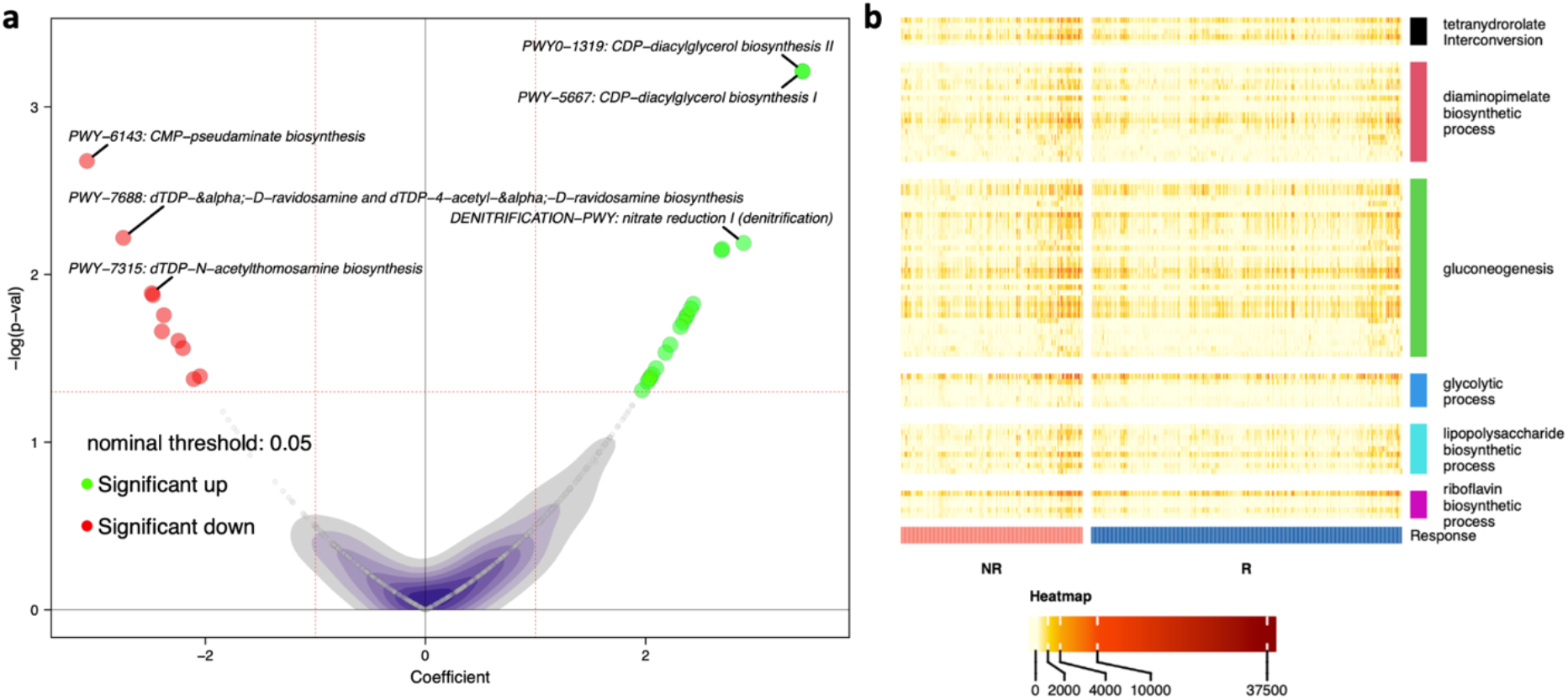
Gene families and pathways associated with immunotherapy response in melanoma. **a**, Pathway Analysis: Pathway enrichment analysis identifying key biological pathways involved in cancer immunotherapy using HUMAnN result. Enrichment scores are plotted, emphasizing pathways like immune response regulation, antigen presentation, and T-cell activation. Significant pathways (p < 0.05, adjusted for multiple comparisons) are annotated for clarity. **b**, Gene Family Analysis: The heatmap reveals distinct metabolic differences between responders (R) and non-responders (NR) to therapy, with certain pathways showing higher activity in each group. Visualization of gene family enrichment across the datasets in the meta-analysis using omePath result. The x-axis displays the gene families, and the y-axis represents the enrichment scores (with significance thresholds denoted by dashed lines). Gene families with strong associations to immune-related functions are highlighted.

The gene families heatmap demonstrated the most metabolic processes (**Fig. 4b**), including tetrahydrofolate interconversion, diaminopimelate biosynthetic process, gluconeogenesis, glycolytic process, lipopolysaccharide biosynthetic process, and riboflavin biosynthetic process. Processes like the glycolytic process and tetrahydrofolate interconversion show higher activity in responders, and processes like gluconeogenesis and lipopolysaccharide biosynthetic process appear to have higher activity in non-responders.

To explore microbial functions beyond pathways and gene families, we analyzed biosynthetic gene clusters (BGCs), which produce specialized metabolites that can influence both microbial interactions and host responses. Across all seven studies, we found that RiPPs, one of the BGC classes, are more abundant in patients who responded well to ICI treatment. At the family level, *Enterobacteriaceae* is also positively linked to treatment response (**Fig. 5**).

**Figure 5:**
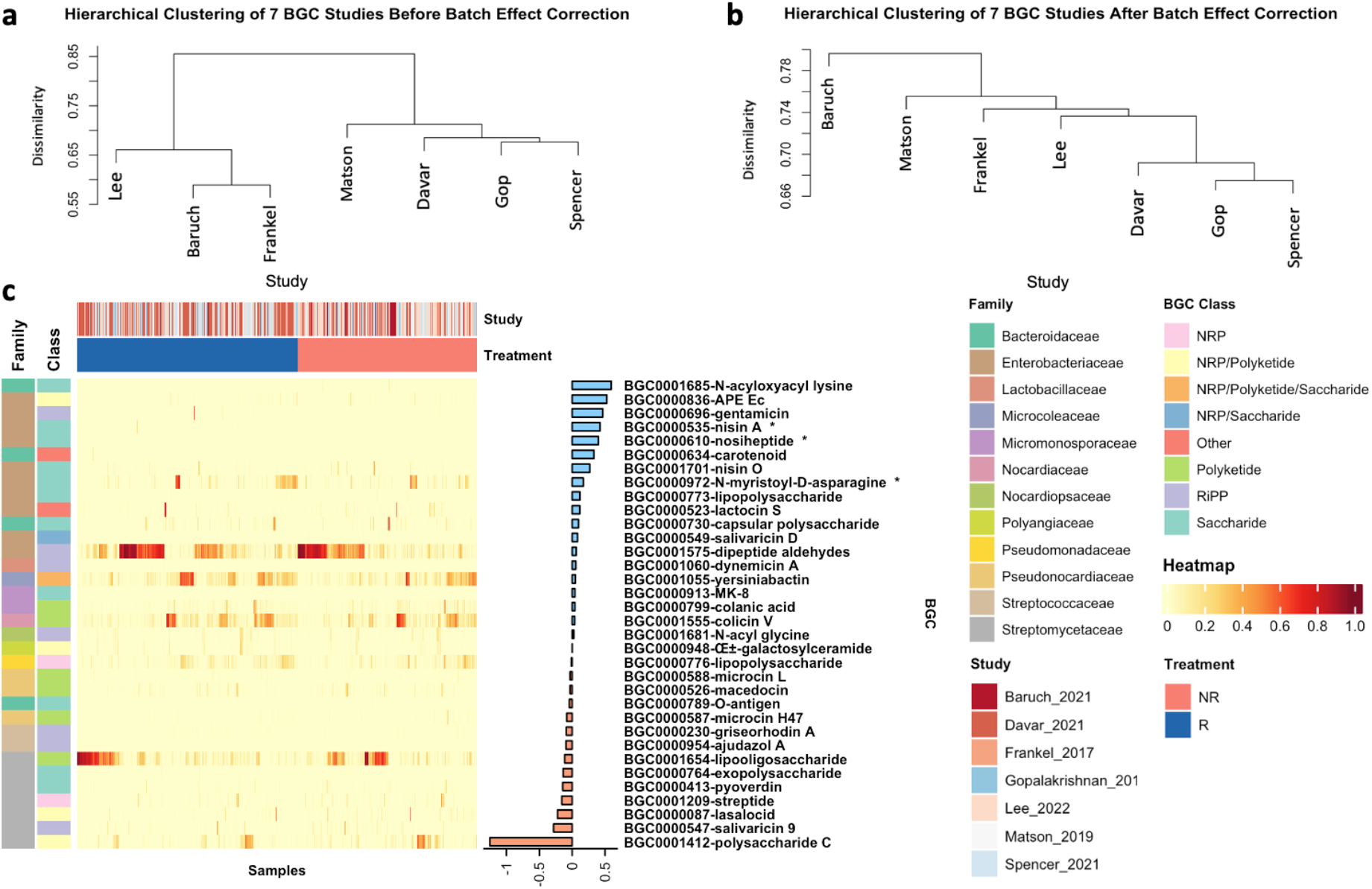
Differences in BGC abundances between responders and nonresponders to ICI therapy in seven studies. **a**, Hierarchical clustering of seven BGC studies before batch effect correction. Before the correction, studies cluster separately, indicating strong batch effects, indicating strong batch effects that contribute to differences in BGC composition across studies. **b**, Hierarchical clustering of seven BGC studies after batch effect correction. Clustering of the same studies after batch effect correction shows improved alignment, with all studies clustering into a single group. Unlike the taxonomy-based analysis, where only a subset of studies could be analyzed together after batch effect correction, the BGC analysis demonstrates that correction was effective across all seven studies, allowing for downstream analyses to include the entire dataset. **c**, The heatmap presents the relative abundance of biosynthetic gene clusters (BGCs) in responders (R, blue) and nonresponders (NR, red) from seven melanoma-associated microbiome cohorts (Baruch et al., 2021; Davar et al., 2021; Frankel et al., 2017; Lee et al., 2022; Matson et al., 2019; Spencer et al., 2021; Gopalakrishnan et al., 2019). Each column represents an individual sample, while each row represents a specific BGC. The color intensity reflects the relative abundance of each BGC, with darker colors indicating higher abundances. The bars at the top indicate the study batch (in shades of blue and red) and treatment response (blue for responders, red for nonresponders). The left sidebar categorizes BGCs by their microbial family (e.g., Bacteroidaceae, Enterobacteriaceae, Streptomycetaceae) and BGC class (e.g., RiPP, NRPs, Polyketide, Saccharide). The bar plot on the right shows the effect size and direction of differences between responders and nonresponders for each BGC, with positive values indicating enrichment in responders and negative values indicating enrichment in nonresponders. Notably, RiPPs were more abundant in responders, supporting their potential role in modulating ICI efficacy.

## Discussion

This study presents a comprehensive meta-analysis that integrates microbial taxonomic profiling, biosynthetic gene cluster characterization, and pathway analysis to elucidate the gut microbiome’s role in modulating ICI responses in advanced melanoma. While numerous individual studies have investigated the effects of PD-1 blockade, fecal microbiota transplantation, and dietary interventions on melanoma treatment, systematic meta-analyses comparing microbiome composition and functional pathways between ICI responders and non-responders remain limited. Notably, this is the first study to analyze BGCs in the context of a microbiome meta-analysis of ICI response, offering novel insights into the functional potential of gut microbial communities in shaping immunotherapy outcomes. By addressing these gaps, our study enhances the understanding of microbiome-mediated immunotherapy modulation and underscores the critical role of gut microbiota in influencing immunotherapy efficacy, highlighting specific microbial species, gene families, and pathways associated with treatment outcomes.

A major highlight of our findings is the identification of *Faecalibacterium* SGB15346 as a robust biomarker for positive ICI responses. This species demonstrated consistent enrichment in responders across multiple datasets, even after rigorous batch effect correction and inter-study variability adjustments. *Faecalibacterium* SGB15346 and *GGB9534* SGB14937 emerged as potential mediators of enhanced immune responses, reinforcing prior evidence that microbiome composition influences immunotherapy outcomes. Among the BGCs, RiPPs (ribosomally synthesized and post-translationally modified peptides) show a positive correlation with the responders and are highly abundant in the responders. Apart from antibacterial activities, many RiPPs have also been shown to have anticancer properties^28^ (**Fig. 5c**). These findings underscore the potential of microbiome-based biomarkers in clinical decision-making and personalized treatment strategies.

Batch effect correction played a critical role in ensuring the reliability of our findings. By implementing state-of-the-art correction techniques, we identified consistent microbial signatures across studies, mitigating one of the primary limitations in microbiome meta-analyses. However, residual clustering suggests that unaccounted confounders, such as medication history and dietary habits, may still influence microbial composition and should be explored in future studies.

Our pathway enrichment analysis revealed the critical functional roles of microbial communities in shaping immunotherapy responses. Notably, pathways involved in SCFA biosynthesis, antigen processing, and T-cell activation were significantly enriched in responders. SCFAs, produced by gut commensals such as *Faecalibacterium* spp., are well-documented for their ability to enhance regulatory T-cell activity and promote an anti-inflammatory microenvironment ^29–31^. These findings provide mechanistic insights into how the gut microbiota modulates the tumor-immune axis, offering potential targets for therapeutic interventions.

The clinical implications of our findings extend beyond biomarker discovery. Emerging therapeutic strategies, such as fecal microbiota transplantation (FMT) and dietary interventions, hold promise in augmenting ICI efficacy by reshaping the gut microbiota ^32,33^. The success of FMT in overcoming PD-1 resistance in recent trials highlights its potential as a complementary therapy. Similarly, our findings support the role of dietary fiber in fostering a microbiome conducive to positive treatment outcomes, suggesting that personalized dietary modifications could enhance immunotherapy efficacy.

Despite these promising findings, our study has certain limitations. Residual batch effects, inherent biological variability, and differences in study design present ongoing challenges in microbiome meta-analysis. Moreover, while our analysis identifies strong correlations, it does not establish causality. Future research should prioritize longitudinal studies and mechanistic experiments to elucidate causal relationships between specific microbial features and immunotherapy responses. Integrating multi-omics data, including metabolomics and transcriptomics, could further enhance our understanding of these complex interactions.

The choice of microbial profiling tools and reference databases significantly influences microbiome analysis outcomes. Tools such as MetaPhlAn 2 and MetaPhlAn 4 rely on different reference genome sets, taxonomic classification levels, and microbial marker selections. These variations can lead to discrepancies in microbial abundance estimations, with some microbes detected by one version but not another or quantified at different levels of accuracy (**Extended Data Fig. 1**). Additionally, frequent database updates incorporating newly discovered taxa and revised taxonomic classifications may introduce biases and inconsistencies, underscoring the need for standardized microbial profiling methodologies in future studies.

In summary, our meta-analysis underscores the pivotal role of the gut microbiome in shaping ICI responses in melanoma. By identifying key microbial species, functional pathways, and gene families associated with treatment outcomes, this study lays the groundwork for microbiome-targeted strategies to optimize immunotherapy. Moving forward, standardized protocols, larger cohorts, and integrated multi-omics approaches will be essential to fully harness the therapeutic potential of the gut microbiota in oncology.

## Methods

### Definition of response to therapy

The assessment of immune checkpoint inhibitor (ICI) efficacy adhered to the RECIST v1.1 criteria. Based on radiographic evaluations, patients were stratified into two groups: those exhibiting a response (characterized as a complete response [CR], partial response [PR], or stable disease [SD]) and those demonstrating nonresponse (progressive disease [PD]). The primary clinical endpoints identified for this study were the overall response rate (ORR) and progression-free survival (PFS), with the latter being defined as the interval commencing with the initial administration of an ICI and culminating at the onset of either disease progression or death from any cause.

### Data collection

We downloaded metagenomic data from four publicly available datasets (GopalakrishnanV_2018, MatsonV_2018, FrankelAE_2017, SpencerCN_2021, BaruchEN_2021, DavarD_2021, and LeeKA_2022) through the Sequence Read Archive using the study accession numbers PRJEB22893^9^, PRJNA399742^8^, PRJNA397906^10^, PRJNA770295^13^, PRJNA678737^12^, PRJNA672867^11^, and PRJEB43119^6^. We excluded any samples taken after the start of ICI therapy, non-metagenomic samples, and nonfecal samples. We classified patients into responder and nonresponder groups according to RECIST 1.1 criteria^34^; patients with complete or partial response, as well as stable disease at first evaluation, were classified as responders, whereas patients with PD were classified as nonresponders.

### Quality control, microbiome taxonomics profiling

For quality control purposes, we subjected the whole metagenomic shotgun sequencing reads from these samples to rigorous filtering. This process was executed using KneadData software^35^, which was applied to the raw reads against the GRCh38 and T2T-CHM13^36^ human read databases. This step resulted in the generation of trimmed reads, effectively cleaning human-originated contamination. Run *fastp* to get the quality control report^37^. Subsequently, the purified reads were processed through MetaPhlAn 4, utilizing the mpa_vJun23_CHOCOPhlAnSGB_202307 database, to classify and determine their relative abundance^18^.

### Pathway and gene family identification

To investigate the molecular underpinnings of cancer immunotherapy, we performed a comprehensive analysis of gene families and biological pathways using HUMAnN3 and omePath pipelines. These methods allowed us to identify key gene families and pathways contributing to immune modulation and therapeutic responses^38,39^. Pathway abundance profiles were generated using the HUMAnN3 (The HMP Unified Metabolic Analysis Network) tool, a computational framework designed to efficiently and accurately quantify microbial communities’ functional and metabolic potential from metagenomic sequencing data. HUMAnN3 utilizes a tiered approach that aligns reads against the UniRef90 database to identify microbial species and map them to annotated pathways. omePath used the uniref90s features from the HUMAnN3 result to get the Gene Ontology (GO) terms.

### Biosynthetic gene cluster profiling

In addition to identifying species and pathways to determine the presence of specific organisms, we aimed to understand their potential functional capabilities, particularly in producing bioactive compounds. Such insights reveal the chemical diversity and potential pharmacological applications within the microbial community. To achieve this, we ran antiSMASH (ANTIbiotics & Secondary Metabolite Analysis SHell) 7.0, a microbial genome mining tool for BGC identification and analysis^40^ and also employed BGCLens, a computational tool specifically designed for the accurate identification and quantification of the BGC^41^. BGCLens utilizes a Bayesian reassignment model to enable precise probabilistic inference based on MIBiG database 3.0^42^, accounting for sequencing uncertainty and improving reading assignment to BGC. This robust approach ensures reliable detection and quantification of BGCs, even within the complexity of diverse microbial community structures. This robust approach ensures reliable detection and quantification of BGCs, even within the complexity of diverse microbial community structures. This provides high-resolution insights into the functional potential of microbial communities, paving the way for understanding their chemical and pharmacological diversity.

### Beta diversity analysis and clustering

Since some studies used different DNA extraction toolkits, we need to investigate beta diversity and the similarity between studies, we calculated Bray-Curtis dissimilarity using the *‘vegdist’* function from the vegan package on a transposed species abundance matrix^43^. Study labels were extracted from sample identifiers to group samples by study. Pairwise mean dissimilarities were computed both within and between studies, with the results stored in a symmetric distance matrix. Hierarchical clustering was then performed on the study-level distance matrix using Ward’s method to minimize variance within clusters. The resulting dendrogram visualized the relationships among studies based on their beta diversity, highlighting ecological similarity in species composition.

### Batch effect correction and meta-analysis

To minimize these batch effects, we implemented a standardized correction protocol targeting taxonomy, BGCs, and pathway profiling data. The effectiveness of this correction was evaluated by calculating *R*^2^ values, which quantify the proportion of variance in the dependent variable that is explained by the independent variables. In the context of batch effect correction, *R*^2^ serves as a key metric, reflecting the extent to which unwanted technical variation has been mitigated. By comparing *R*^2^ values before and after correction, we assessed improvements in data quality and the reduction of batch-associated biases^44,45^.

Data from multiple studies were integrated using batch effect removal and meta-analysis. Batch correction was performed with the ‘adjust_batch’ function from the MMUPHin R package^46^, ensuring consistent interpretation across datasets. Statistical significance and the explanatory power of variables were assessed using permutational multivariate analysis of variance (PERMANOVA), executed via the ‘adonis’ function in the vegan R package. For further analysis, *R*^2^ values were determined to evaluate both statistical significance and the variance explained by the models. To explore associations between response variables and feature abundances while accounting for batch effects, we conducted a meta-analysis using the ‘lm_meta’ function, derived from MaAsLin2^47^. This analysis employed a compound Poisson linear model (CPLM^48^) with the response variable designated as the exposure and the dataset identifier included as both a batch variable and a random effect.

The input to the CPLM consisted of a batch-adjusted feature abundance matrix, with metadata specifying the response variable and dataset identifiers. The model did not incorporate additional transformations, normalization, or standardization and used “Non-Response” as the reference category for the response variable.

## Data availability

The seven publicly available datasets include metagenomes, and main metadata relevant to the analyses are deposited in the European Nucleotide Archive under accession numbers PRJEB22893, PRJNA399742, PRJNA397906, PRJNA770295, PRJNA678737, PRJNA672867, All MetaPhlAn 4 and HUMAnN 3 profiles are available within the latest version of curatedMetagenomicData (https://bioconductor.org/packages/curatedMetagenomicData/).

## Code availability

The code used to analyze data and generate figures from this project is available at https://github.com/omicsEye/Cancer-Microbiome

## Acknowledgments

This work was partially supported by the National Science Foundation grant number 2109688 to AR. We acknowledge Dr. Jia (John) Kang (Merck & Co., Inc., Rahway, NJ, USA) for providing initial feedback on an earlier version of the meta-analysis.

## Author information

### Authors and Affiliations

**Computational Biology Institute, Department of Biostatistics and Bioinformatics, Milken Institute School of Public Health, The George Washington University, Washington, DC, USA**

Xinyang Zhang, Ali Rahnavard

**Division of Biostatistics, Department of Population Health Sciences, Weill Cornell Medicine, Cornell University, New York, NY, USA**

**Department of Statistics and Data Science, Cornell University, Ithaca, NY, USA**

Himel Mallick

## Authors Contribution

A.R. and H.M. conceptualized and designed the study, providing overall project oversight. X.Z. conducted data preparation, performed bioinformatic analyses, created visualizations, and drafted the manuscript. All authors contributed to the interpretation of the results, provided critical feedback, and participated in manuscript writing.

## Ethics declarations

### Conflicts of Interest

No authors have any conflicts of interest to declare.

## Extended data

**Extended Data Figure 1:**
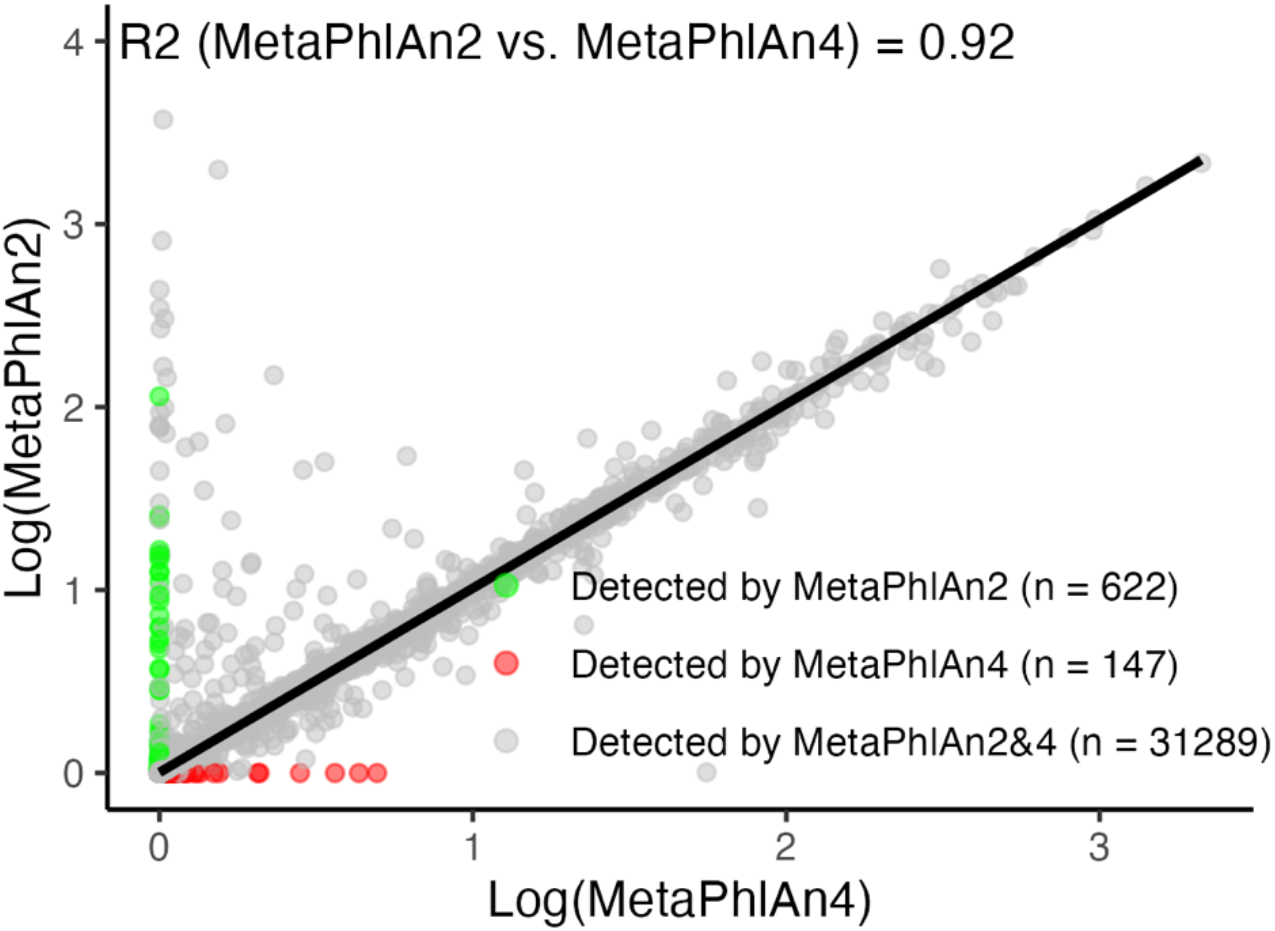
Scatterplot depicting the correlation between microbial abundances as determined by two different versions of the MetaPhlAn tool (MetaPhlAn 2 vs. MetaPhlAn 4). We ran the ‘Frankel’ dataset on two different databases to make the comparison. The logarithmic transformation of the abundance data is plotted on both axes, facilitating a comparison of the two analytical methods. A strong positive correlation (R^2^ = 0.92) is evident, indicating a high level of agreement between the datasets generated by each version. The blue line represents the line of best fit, demonstrating the consistency across the two profiling methodologies. Species with zero read counts in MetaPhlAn 2 and positive values in MetaPhlAn 4 can reflect new species detected in MetaPhlAn 4. In contrast, those with zero in MetaPhlAn 4 and positive values in MetaPhlAn 2 could be false positives in MetaPhlAn 2.

**Extended Data Figure 2:**
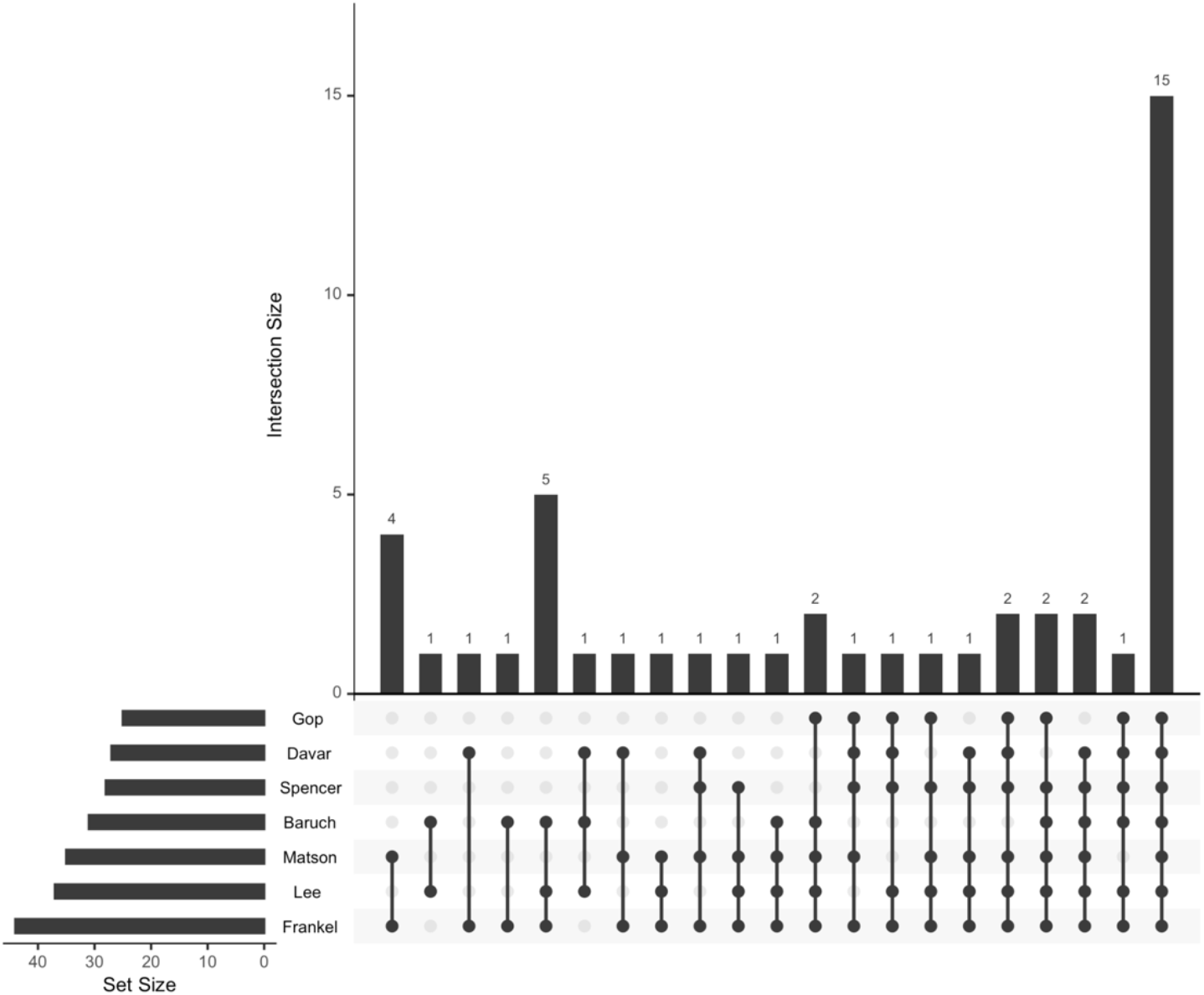
UpSet plot shows the intersection among studies species. UpSet plot shows the intersections of significant microbial species across seven studies. It illustrates the shared and unique species among the datasets labeled Gop, Baruch, Matson, Davar, Frankel, Spencer, and Lee. The bar graph at the top shows the size of each intersection, indicating the number of significant species, with p-val<0.05, common to each combination of datasets. The horizontal bars on the left represent the total number of species identified in each study. The connected dots below the bars indicate the specific datasets involved in each intersection. Notably, the largest intersection involves species shared by all seven datasets, with additional significant intersections involving various combinations of two or more datasets.

**Extended Data Figure 3:**
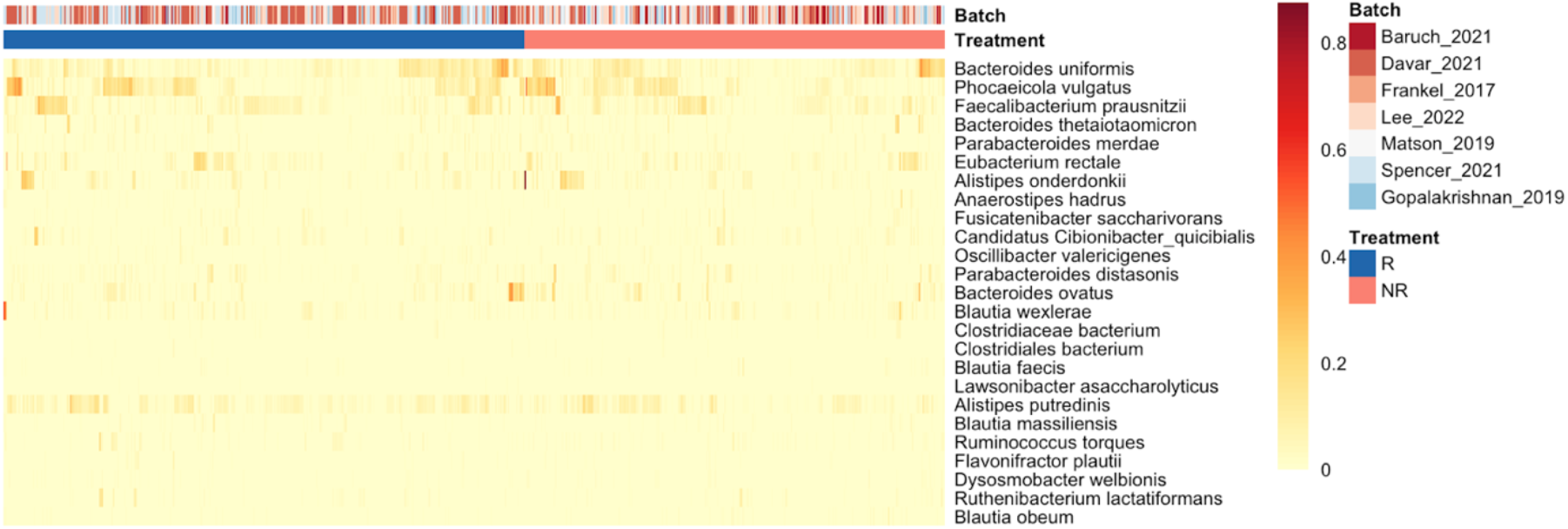
Differences in microbial abundances between responders and nonresponders to ICI therapy in seven studies. The heatmap shows the relative abundance of microbial species across responders (R, blue) and nonresponders (NR, red) from seven melanoma-associated microbiome cohorts (Baruch et al., 2021; Davar et al., 2021; Frankel et al., 2017; Lee et al., 2022; Matson et al., 2019; Spencer et al., 2021; Gopalakrishnan et al., 2019). Each column represents an individual sample, and each row represents a microbial species. The color intensity indicates the relative abundance of each species, with darker colors representing higher abundances. The top bars indicate the study batch (in shades of red) and treatment response (blue for responders and red for nonresponders). Species such as *Faecalibacterium prausnitzii, Bacteroides uniformis*, and *Eubacterium rectale* show higher abundances in responders, highlighting potential microbial signatures associated with ICI treatment efficacy.

**Extended Data Table 1:**
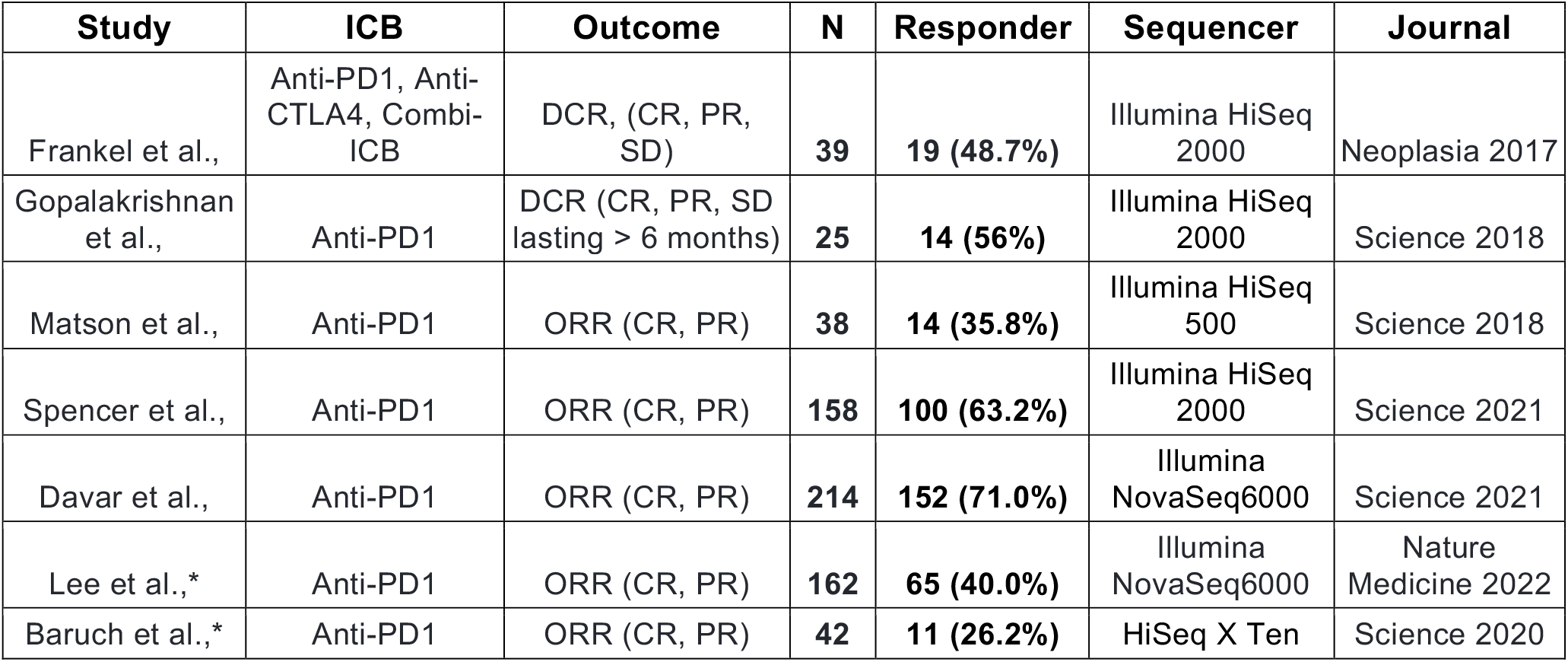
Summary of the studies used for the meta-analysis. Total =678; Abbreviations: CR, Complete response, PR; Partial response; DCR, Disease control rate; MM, metastatic melanoma; MGS, metagenomic sequencing; OS, Overall survival; ORR, Objective response rate; PFS, Progression-free survival; SD, Stable disease; ICB, immune checkpoint blockade; Study name with asterisk(*) did not share the most similarity with other studies on taxonomy profiling result.

| Study | ICB | Outcome | N | Responder | Sequencer | Journal |
| --- | --- | --- | --- | --- | --- | --- |
| Frankel et al., | Anti-PD1, Anti-CTLA4, Combi-ICB | DCR, (CR, PR, SD) | 39 | 19 (48.7%) | Illumina HiSeq 2000 | Neoplasia 2017 |
| Gopalakrishnan et al., | Anti-PD1 | DCR (CR, PR, SD lasting > 6 months) | 25 | 14 (56%) | Illumina HiSeq 2000 | Science 2018 |
| Matson et al., | Anti-PD1 | ORR (CR, PR) | 38 | 14 (35.8%) | Illumina HiSeq 500 | Science 2018 |
| Spencer et al., | Anti-PD1 | ORR (CR, PR) | 158 | 100 (63.2%) | Illumina HiSeq 2000 | Science 2021 |
| Davar et al., | Anti-PD1 | ORR (CR, PR) | 214 | 152 (71.0%) | Illumina NovaSeq6000 | Science 2021 |
| Lee et al.,* | Anti-PD1 | ORR (CR, PR) | 162 | 65 (40.0%) | Illumina NovaSeq6000 | Nature Medicine 2022 |
| Baruch et al.,* | Anti-PD1 | ORR (CR, PR) | 42 | 11 (26.2%) | HiSeq X Ten | Science 2020 |

## Rights and permissions

### Open Access

This article is licensed under a Creative Commons Attribution 4.0 International License, which permits use, sharing, adaptation, distribution and reproduction in any medium or format, as long as you give appropriate credit to the original author(s) and the source, provide a link to the Creative Commons license, and indicate if changes were made. The images or other third-party material in this article are included in the article’s Creative Commons license unless indicated otherwise in a credit line to the material. If material is not included in the article’s Creative Commons license and your intended use is not permitted by statutory regulation or exceeds the permitted use, you will need to obtain permission directly from the copyright holder. To view a copy of this license, visit http://creativecommons.org/licenses/by/4.0/.

## Notes

https://github.com/omicsEye/Cancer-Microbiome

### Competing Interest Statement

The authors have declared no competing interest.

## References

1. Schadendorf, D. et al. Melanoma. Lancet 392, 971–984 (2018).

2. Routy, B. et al. Melanoma and microbiota: Current understanding and future directions. Cancer Cell 42, 16–34 (2024).

3. van Not, O. J. et al. Long-Term Survival in Patients With Advanced Melanoma. JAMA Netw Open 7, e2426641 (2024).

4. Boutros, A. et al. The treatment of advanced melanoma: Current approaches and new challenges. Crit Rev Oncol Hematol 196, 104276 (2024).

5. Routy, B. et al. Gut microbiome influences efficacy of PD-1-based immunotherapy against epithelial tumors. Science 359, 91–97 (2018).

6. Lee, K. A. et al. Cross-cohort gut microbiome associations with immune checkpoint inhibitor response in advanced melanoma. Nat. Med. 28, 535–544 (2022).

7. Villemin, C. et al. The heightened importance of the microbiome in cancer immunotherapy. Trends Immunol 44, 44–59 (2023).

8. Matson, V. et al. The commensal microbiome is associated with anti-PD-1 efficacy in metastatic melanoma patients. Science 359, 104–108 (2018).

9. Gopalakrishnan, V. et al. Gut microbiome modulates response to anti-PD-1 immunotherapy in melanoma patients. Science 359, 97–103 (2018).

10. Frankel, A. E. et al. Metagenomic Shotgun Sequencing and Unbiased Metabolomic Profiling Identify Specific Human Gut Microbiota and Metabolites Associated with Immune Checkpoint Therapy Efficacy in Melanoma Patients. Neoplasia 19, 848–855 (2017).

11. Davar, D. et al. Fecal microbiota transplant overcomes resistance to anti-PD-1 therapy in melanoma patients. Science 371, 595–602 (2021).

12. Baruch, E. N. et al. Fecal microbiota transplant promotes response in immunotherapy-refractory melanoma patients. Science 371, 602–609 (2021).

13. Spencer, C. N. et al. Dietary fiber and probiotics influence the gut microbiome and melanoma immunotherapy response. Science 374, 1632–1640 (2021).

14. Limeta, A., Ji, B., Levin, M., Gatto, F. & Nielsen, J. Meta-analysis of the gut microbiota in predicting response to cancer immunotherapy in metastatic melanoma. JCI Insight 5, (2020).

15. Gunjur, A. et al. A gut microbial signature for combination immune checkpoint blockade across cancer types. Nat Med 30, 797–809 (2024).

16. Olekhnovich, E. I. et al. Consistent Stool Metagenomic Biomarkers Associated with the Response To Melanoma Immunotherapy. mSystems 8, e0102322 (2023).

17. Truong, D. T. et al. MetaPhlAn2 for enhanced metagenomic taxonomic profiling. Nat Methods 12, 902–903 (2015).

18. Blanco-Míguez, A. et al. Extending and improving metagenomic taxonomic profiling with uncharacterized species using MetaPhlAn 4. Nat Biotechnol 41, 1633–1644 (2023).

19. Nouws, S. et al. Impact of DNA extraction on whole genome sequencing analysis for characterization and relatedness of Shiga toxin-producing Escherichia coli isolates. Sci Rep 10, 14649 (2020).

20. Hart, M. L., Meyer, A., Johnson, P. J. & Ericsson, A. C. Comparative Evaluation of DNA Extraction Methods from Feces of Multiple Host Species for Downstream Next-Generation Sequencing. PLoS One 10, e0143334 (2015).

21. Wang, Y. & Lê Cao, K.-A. PLSDA-batch: a multivariate framework to correct for batch effects in microbiome data. Brief Bioinform 24, (2023).

22. Thomas, P. D. et al. PANTHER: Making genome-scale phylogenetics accessible to all. Protein Sci 31, 8–22 (2022).

23. Tagore, M. et al. GABA Regulates Electrical Activity and Tumor Initiation in Melanoma. Cancer Discov 13, 2270–2291 (2023).

24. Ceol, C. J. Microenvironmental GABA Signaling Regulates Melanomagenesis through Reciprocal Melanoma-Keratinocyte Communication. Cancer Discov 13, 2128–2130 (2023).

25. Molagoda, I. M. N. et al. Gamma-Aminobutyric Acid (GABA) Inhibits α-Melanocyte-Stimulating Hormone-Induced Melanogenesis through GABA and GABA Receptors. Int J Mol Sci 22, (2021).

26. Oikari, S. et al. UDP-sugar accumulation drives hyaluronan synthesis in breast cancer. Matrix Biol 67, 63–74 (2018).

27. Hyaluronan in human malignancies. Experimental Cell Research 317, 383–391 (2011).

28. RiPP antibiotics: biosynthesis and engineering potential. Current Opinion in Microbiology 45, 61–69 (2018).

29. Fusco, W. et al. Short-Chain Fatty-Acid-Producing Bacteria: Key Components of the Human Gut Microbiota. Nutrients 15, (2023).

30. Ney, L.-M. et al. Short chain fatty acids: key regulators of the local and systemic immune response in inflammatory diseases and infections. Open Biol 13, 230014 (2023).

31. Hersi, F. et al. Cancer immunotherapy resistance: The impact of microbiome-derived short-chain fatty acids and other emerging metabolites. Life Sci 300, 120573 (2022).

32. Zhang, J., Wu, K., Shi, C. & Li, G. Cancer Immunotherapy: Fecal Microbiota Transplantation Brings Light. Curr Treat Options Oncol 23, 1777–1792 (2022).

33. Yang, Y. et al. Fecal microbiota transplantation: no longer cinderella in tumour immunotherapy. EBioMedicine 100, 104967 (2024).

34. Eisenhauer, E. A. et al. New response evaluation criteria in solid tumours: revised RECIST guideline (version 1.1). Eur J Cancer 45, 228–247 (2009).

35. GitHub - biobakery/kneaddata: Quality control tool on metagenomic and metatranscriptomic sequencing data, especially data from microbiome experiments. GitHub https://github.com/biobakery/kneaddata.

36. Nurk, S. et al. The complete sequence of a human genome. Science (2022) doi:10.1126/science.abj6987.

37. Chen, S., Zhou, Y., Chen, Y. & Gu, J. fastp: an ultra-fast all-in-one FASTQ preprocessor. Bioinformatics 34, i884–i890 (2018).

38. GitHub - omicsEye/omePath: omePath: Omics Pathway Enrichment Analysis. GitHub https://github.com/omicsEye/omePath.

39. Beghini, F. et al. Integrating taxonomic, functional, and strain-level profiling of diverse microbial communities with bioBakery 3. Elife 10, (2021).

40. Blin, K. et al. antiSMASH 7.0: new and improved predictions for detection, regulation, chemical structures and visualisation. Nucleic Acids Res 51, W46–W50 (2023).

41. GitHub - omicsEye/BGCLens. GitHub https://github.com/omicsEye/BGCLens.

42. Terlouw, B. R. et al. MIBiG 3.0: a community-driven effort to annotate experimentally validated biosynthetic gene clusters. Nucleic Acids Res. 51, D603–D610 (2022).

43. Sherwin, W. B. Bray-Curtis (AFD) differentiation in molecular ecology: Forecasting, an adjustment (), and comparative performance in selection detection. Ecol Evol 12, e9176 (2022).

44. Tran, H. T. N. et al. A benchmark of batch-effect correction methods for single-cell RNA sequencing data. Genome Biol 21, 12 (2020).

45. Yu, Y., Mai, Y., Zheng, Y. & Shi, L. Assessing and mitigating batch effects in large-scale omics studies. Genome Biol 25, 254 (2024).

46. Ma, S. et al. Population structure discovery in meta-analyzed microbial communities and inflammatory bowel disease using MMUPHin. Genome Biol. 23, 1–31 (2022).

47. Mallick, H. et al. Multivariable association discovery in population-scale meta-omics studies. PLoS Comput. Biol. 17, e1009442 (2021).

48. Mallick, H. et al. Differential expression of single-cell RNA-seq data using Tweedie models. Stat Med 41, 3492–3510 (2022).

